# Computational search of hybrid human/ SARS-CoV-2 dsRNA reveals unique viral sequences that diverge from those of other coronavirus strains

**DOI:** 10.1101/2020.04.08.031856

**Authors:** Claude Pasquier, Alain Robichon

## Abstract

The role of the RNAi/Dicer/Ago system to degrade RNA viruses has been elusive in mammals, which prompted authors to think that interferon (IFN) synthesis is essential in this clade relegating the RNAi defense strategy against viral infection as accessory function. We explore the theoretical possibilities that RNAi triggered by SARS-CoV-2 might degrade some host transcripts in the opposite direction although this hypothesis seems counter intuitive. SARS-CoV-2 genome was therefore computational searched for exact intra pairing within the viral RNA and also hybrid exact pairing with human transcriptome over a minimum 20 bases length. Minimal segments of 20 bases length of SARS-CoV-2 RNA were found based on the theoretical matching with existing complementary strands in the human host transcriptome. Few human genes potentially annealing with SARS-CoV-2 RNA, among them mitochondrial deubiquitinase USP30, a subunit of ubiquitin protein ligase complex FBXO21 along with two long coding RNAs were retrieved. The hypothesis that viral originated RNAi might mediate degradation of messengers of the host transcriptome was corroborated by clinical observation and phylogenetic comparative analysis indicating a strong specificity of these hybrid pairing sequences for both SARS-CoV-2 and human genomes.

## Introduction

Two restraint scale epidemics occurred already in the recent past caused by two coronaviruses: the severe acute respiratory syndrome (SARS) and Middle East respiratory syndrome (MERS) [1,2]. The third recent new viral outbreak causing a fast propagating pandemic is provoked by a new coronavirus, named SARS-CoV-2, that exhibits terrifying spread out of control. Metagenomic sequencing along with the phylogenetic analysis of the virus from a sample of broncho-alveolar lavage fluid of an infected patient have been performed [1,2]. The high homology of the predicted protein of the RBD domain of the spike protein with the precedent SARS-CoV hints that the human angiotensin-converting enzyme 2 (ACE2) acts as a receptor for fixation and entry to human host cells [3,4]. Inside mammal infected host cells, the role of RNAi machinerie to degrade virus is debated. In fungi, plants and invertebrates, the viral dsRNAs are trapped and cleaved by Dicer/Ago machinerie into small interfering RNAs (siRNAs) that degrade the targeted viral messenger RNAs within the RNA-induced silencing complex (RISC) [5]. In mammals/humans the detection of virus-derived small RNAs has been mainly undetectable upon RNA virus infection [6]. The unique Dicer in humans processed long double-stranded RNA (dsRNA) and hairpin dsRNA into small interfering RNAs (siRNAs) and microRNAs respectively but its role as producer of antiviral RNAis is yet far to be admitted by many authors. However, mammalian viruses were successfully eliminated and replication inhibited in cultured host cells via synthetic small interfering RNAs (siRNAs) [6]. In mammals, a powerful and non-sequence-specific synthesis of interferon (IFN) appears the major molecular weapon to fight RNA viruses [7]. This effector response might have overwhelmed the initial RNAi role, which is inferred by the rare accumulation of virus-derived small RNAs (vsRNAs) in few mammal host cells for which any specified functions have been found [7–9]. The question whether RNA virus infection of mammalian cells can trigger an effective and powerful antiviral RNAi weapon remains unknown and little documented as no siRNAs of viral origin were experimentally found in mammal infected cells in contrast with the observations in plants and invertebrates [8,9]. The absence of virus specific siRNAs in mammal cells remains presently a puzzling observation for which an explanation could be the scarcity of intra RNA pairing offered by the virus genomes. In the past weeks we proceeded to a computational search of the SARS-CoV-2 full RNA sequence to explore whether short readings presenting perfect match in reverse complementary strand with RNA candidates from the full human transcriptome might exist. We also examined the possibilities of intra pairing within the full SARS-CoV-2 RNA at a minimum 20 bases length. Comparative studies were conducted with the SARS-CoV, the MERS-CoV along with two non-virulent coronaviruses (HCoV-229E and HCoV-OC43). The purpose of this searching was guided by the hypothesis that hybrid duplex RNA (one strand from human transcriptome, the other from SARS-CoV-2) could be theoretically formed and consequently might lead to degradation of RNA targets not only in SARS-CoV-2 but more importantly, the other way around, in the human transcriptome.

Dicer can trap dsRNAs pairing over a 19-base length with at least a 2 nt overhang at the 3’ ends and can then directly transfer the modified/unmodified siRNA to the Ago2 site [10–12]. This mechanism has been further demonstrated with fluorometric dsRNA [13]. Some paired 19-mers with a dTdT overhang at the 3’ end bind Dicer with high affinity; these sequences are not cleaved by Dicer, are fully transferred to Ago and trigger high-efficiency gene silencing activity [10,11]. Dicer also binds single RNAs at a specific site with high affinity, the function of which is to accelerate hybridization with a partner from the “soup” of RNA metabolism [14]. Authors have also shown that the pairing length can be as short as 16 nt, with a several-base overhang at the 3’ end to fully bind to Dicer and consecutively to activate Ago in the RISC context [10–12]. In this study, a pairing length for computational searching was set minimally to 20 bases, which leads to 24 mers considering the 2nt overhang in both 3’ extremities for Dicer cleavage specificity. Exact duplex at and/or above 20 bases length coming from intra annealing of viral RNA were not found although the possibilities of imperfect or very short pairing length exist. Inversely the computational search of hybrid pairing RNAs has retrieved few human genes related to ubiquitin metabolism and two long non coding RNAs. This observation validate the hypothesis that RISC/Ago might be fueled by diverse source of RNAi, byproducts of the action of Dicer, originated from hybrid and asymmetric RNA duplex, one strand coming from host and the other from the virus.

## Results

### List of human transcripts theoretically duplexing with SARS-CoV-2 RNA

Short segments of SARS-CoV-2 RNAs that are potential sources of siRNA by hybridization with human RNAs were computationally found, which identified the transcripts of 7 coding human genes that are complementary to a segment of SARS-CoV-2 RNA. Results are summarized in Table 1 and the full list of pairing transcript isoforms is shown in Table S1. Among the human genes, we noticed the presence of DNAJC13 (which regulates endosomal membrane trafficking [15]), FBXO21 (a F-box protein that is one of the four subunits of ubiquitin protein ligase complex [16]), FLRT2 (encodes a fibronectin leucine rich transmembrane), ELP4 (encodes an histone acetyltransferase, a subunit associated with RNA polymerase type II), USP31 (an ubiquitin specific peptidase that has been described to activate transcription factor NF-kappaB that stimulates interferon synthesis [17]) and USP30 (a mitochondrial ubiquitin specific peptidase [18]). The segment of USP30 RNA having a reverse complement in SARS-CoV-2 RNA was found exclusively in humans, which highlights the unexpected and intriguing specificity of this coronavirus. The viral pairing sequences are originating from 3 putative open reading frames (orf): orf1a/b, Spike and orf3b. In a recent study, Chan et al., show that there are no remarkable differences between orf1a/b in SARS-CoV-2 with the one in SARS-CoV [19], the major distinction being located in orf3b, Spike and orf8. The authors note that orf3a encodes a completely novel short protein that may have a role in viral pathogenicity. Orf3a also includes the 20-base sequence that potentially targets the ubiquitin specific peptidase 30 (USP30). The complementary sequence segment of USP30 within the SARS-CoV-2 was found only in this coronavirus strain whereas the corresponding USP30 segment was not found in other mammal species.

**Table 1:**
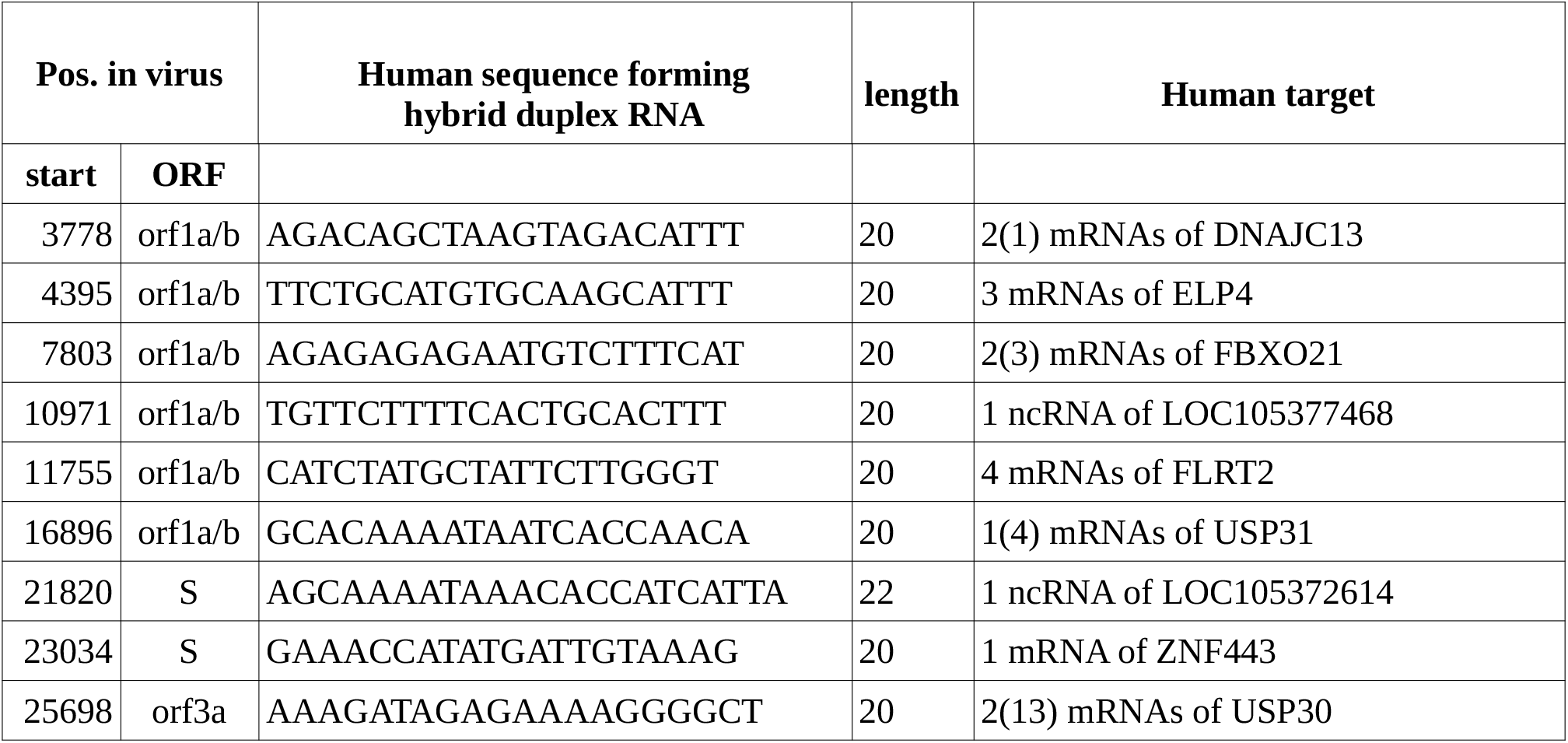
List of hybrid duplex RNA potentially formed with one strand from human transcriptome and the other from SARS-CoV-2. The table shows the position of the sequence in the RNA virus (start position and ORF name), the DNA sequence coding the human RNA and the corresponding features in human. The numbers in the column ‘Human target’ are specified using the format x(y) where x and y are the numbers of known and predicted transcripts respectively.

### List of human transcripts theoretically duplexing with RNA of other coronaviruses

The same type of analysis was conducted with other Coronaviridae for comparison: SARS-CoV (having caused the precedent China pandemic), MERS-CoV and two no virulent strains (HCoV-229E and HCoV-OC43). Results are shown in Tables 2–4 and Table S1. We notice that each strain presents a unique set of sequences of 20 bases length matching in reverse complement with human transcripts and these unique sets target no human genes in common. Remarkably, SARS-CoV mostly targets ncRNAs (Table 2 and Table S1). MERS targets an histone methyltransferase (KMT2C) and a miscRNA included in this gene; a small GTPase of the Rho-subfamily (CDC42); an ubiquitin protein ligase (ARH1); a calcium activated chloride channels (ANO9); a voltage and calcium-sensitive potassium channel (KCNMA1) and finally an adhesion glycoprotein (THBS3) (Table 3 and Table S1). Regarding the no virulent coronavirus (HCoV-229E and HCoV-OC43), few human genes are potentially targeted by viral RNA sequences, among them few no coding RNAs for which a role in defense against viral infection is not documented. The few coding genes do not appear to qualify for GO annotation of cellular signaling susceptible to be altered by the virus infection (see NCBI database for information).

**Table 2:**
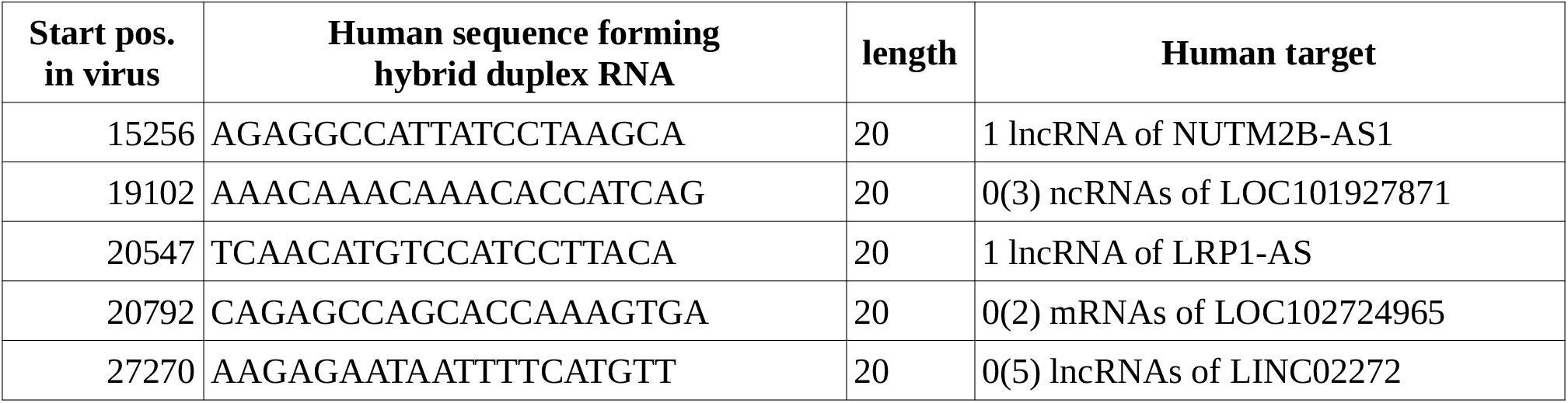
List of hybrid duplex RNA potentially formed with one strand from human transcriptome and the other from SARS-CoV. The table shows the start position of the sequence in the RNA virus, the DNA sequence coding the human RNA and the corresponding features in human. The numbers in the column ‘Human target’ are specified using the format x(y) where x and y are the numbers of known and predicted transcripts respectively.

**Table 3:**
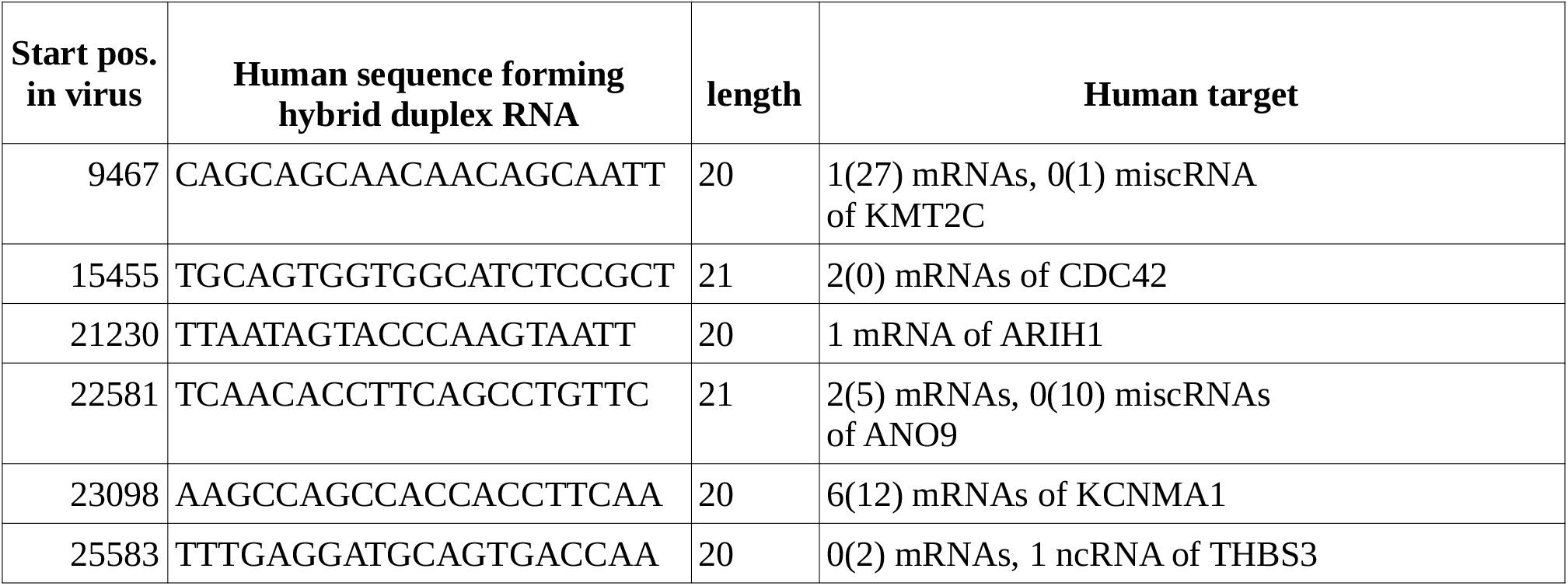
List of hybrid duplex RNA potentially formed with one strand from human transcriptome and the other from MERS-CoV. The table shows the start position of the sequence in the RNA virus, the DNA sequence coding the human RNA and the corresponding features in human. The numbers in the column ‘Human target’ are specified using the format x(y) where x and y are the numbers of known and predicted transcripts respectively.

**Table 4:**
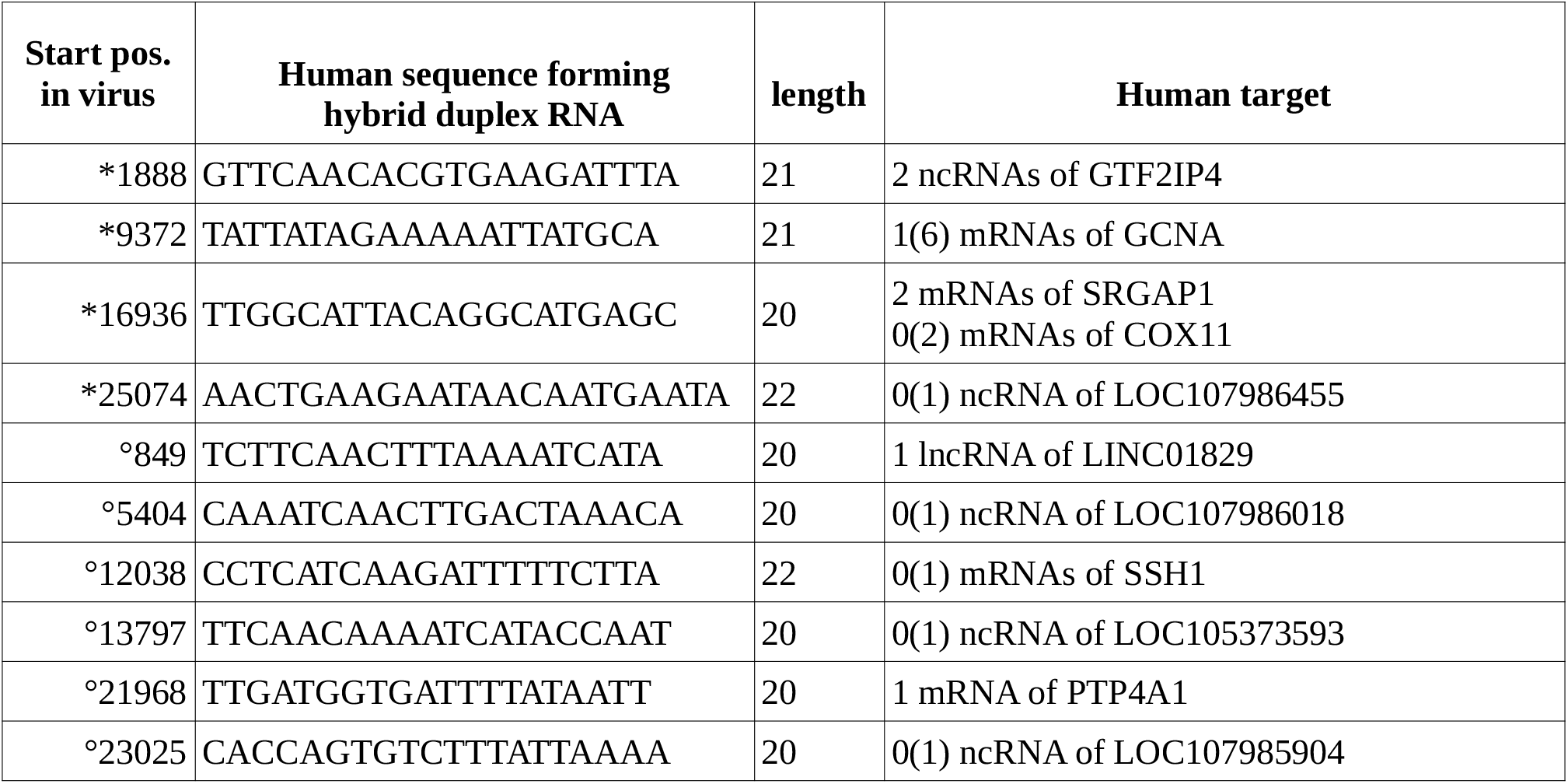
List of hybrid duplex RNA potentially formed with one strand from human transcriptome and the other from HCoV-229E and HCoV-OC43. The 4 first lines (*1888 to 25074) refers to HCoV-229E and the 6 last lines (°849 to 23025) to HCoV-OC43. The table shows the start position of the sequence in the RNA virus, the DNA sequence coding the human RNA and the corresponding features in human.The numbers in the column ‘Human target’ are specified using the format x(y) where x and y are the numbers of known and predicted transcripts respectively.

### Theoretical hybrid human/ SARS-CoV-2 dsRNA are related to ubiquitin pathway and mitochondria

Upon RNA viral infection, the RNA helicases (RIG-I and MDA-5) binds virus-derived RNAs [20–22]. The complexes then translocate at the outer mitochondrial membrane by binding with MAVS (mitochondrial antiviral signaling). The bound MAVS acting as scaffold protein recruits downstream effectors to form a “MAVS signalosome” of which the major function resides in drastically inducing the stimulation of the NF-κB and IRF-3 factors; a molecular scenario well documented in viral infection of host mammal cells [23]. Mitochondria appear to constitute a hub of communication that implicates a cascade of effector molecules recruited for antiviral defense [23]. Regarding the humans genes targeted by SARS-CoV-2, we observe the presence of USP30 a deubiquitinase specific of mitochondria and FBXO21which is a subunit in an ubiquitin protein ligase complex. Remarkably FBXO21 ubiquitin ligase complex has been reported to be required for antiviral innate response [16]. The host ubiquitin system is known to be crucial in innate immunity, the host E3-ubiquitin ligase having clearly antiviral functions [24]. Putatively dsRNAs, hybrid forms with one strand coming from human transcriptome and the other from viral RNA might have the unfortunate role to direct the Dicer/ago/RISC defense system against its own host.

### Philogenetic analysis of the 20 bases length of RNA sequences between coronaviruses strains and between mammal species

We conducted the comparison of the RNA segments within the ortholog genes of numerous vertebrate species with a particular emphasis placed on the class of primates. Results are shown in figure 1 and Supplemental data-Table S2-S8. In summary, one 20 bases sequence (FBXO21) presents 100% homology between human, gorilla and chimpanzee. One other sequence (USP30) is unique to human whereas the sequence (USP31) was found in human, gorilla, chimpanzee and capuchin. All these sequences were found to be phylogenetically distant in other mammal species. Overall the combinatorial setting of the 7 RNA segments in humans was unmatched with another primate species, which highlights that a pattern of gene targets might be unique for humans. The same type of analysis was conducted between different strains of coronaviruses (see figure 2 and Supplemental data-Tables S9-S15). In such case the conservation of the 20 bases sequences are weak with significant divergences. We note that Sars-Cov-2 and the bat coronavirus RaTG13 present 4 sequences on 7 identical or with one base change whereas the other coronaviruses known as causing the same clinical symptoms and physiopathology in humans than Sars-Cov-2 are highly divergent for the 7 sequences.

**Figure 1.**
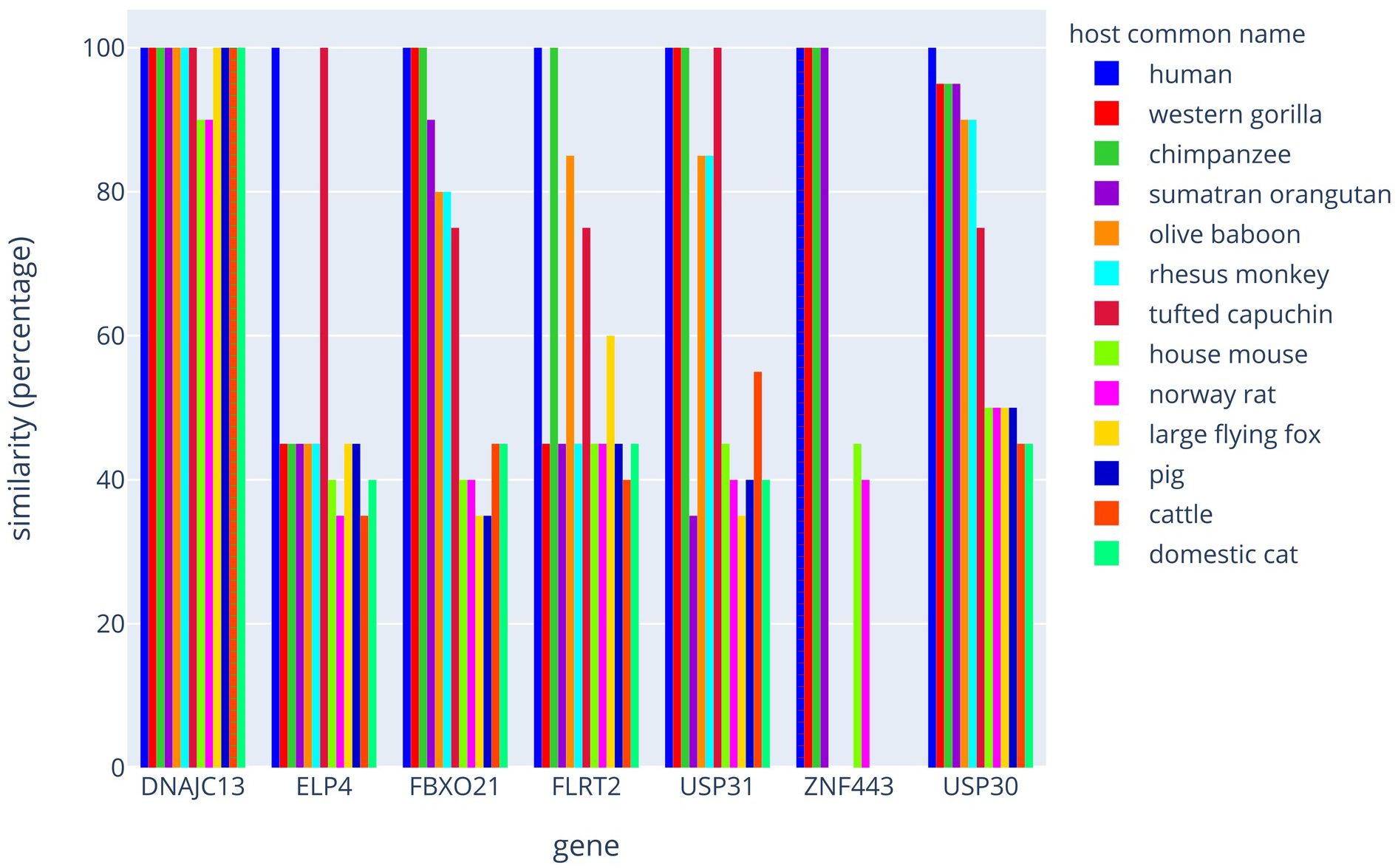
Philogenetic comparison of the human strand of hybrid dsRNAs with other mammals species. The graph represents the comparison of the sequences within the ortholog genes between few mammal species. Results are plotted as a percentage of similarities with the human counterpart.

**Figure 2.**
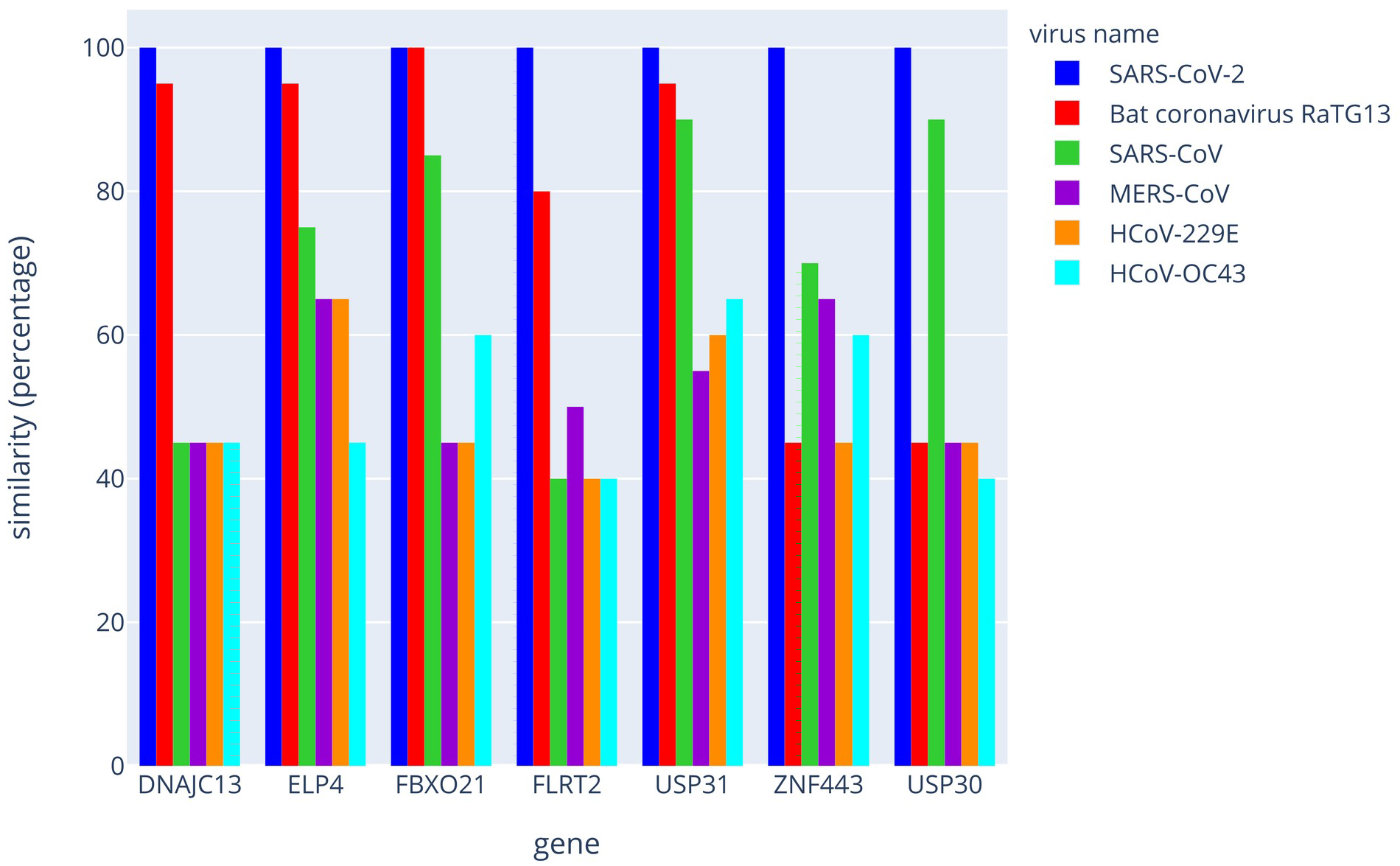
Philogenetic comparison of the SARS-CoV-2 strand of hybrid dsRNAs with other coronaviruses. The graph represents the comparison of the 20 bases sequences between few coronaviruses strains. Results are plotted as a percentage of similarities with the SARS-CoV-2.

## Discussion

Our observations lead to conclusions that i) segments presenting intra or inter RNA pairing above 20 bases length within the 5 coronaviruses RNAs are rare although pairing on very short length and/or pairing with mismatches are frequent, ii) the list of hybrid duplex coranoviruses/human RNAs was significantly more important in SARS-CoV-2 than in the other strains if we consider all the isoforms in the transcriptome, iii) regarding SARS-CoV-2, amazingly hybrid duplexing sequences were found in three important human genes: mitochondrial ubiquitin specific peptidase 30 (USP30), a subunit of ubiquitin ligase complex (FBXO21) and finally ubiquitin specific peptidase 31 (USP31). SARS-CoV-2 might affect mitochondria via ubiquitination alteration, which could be a plausible pathway of interference to explain the dramatic deteriorating conditions of many patients affected by this virus. SARS-CoV-2 might have a “stolen” piece of genetic information from homo sapiens and/or humanoids acquired along the co-evolution history over millions years or alternatively SARS-CoV-2 genome might have randomly drifted allowing the apparition of matching pieces of sequences with human genome; all these genetic innovations could have occurred with *in fine* the catastrophic result which consists to return the human immunity defense system against itself. This argues in favor of the idea that the immune defense Dicer/Ago/RISC system could work against its host upon viral infection. Not only RNA viruses over million years of co-evolution dodge the immune system of hosts but in some cases could trigger a “self destruction” or “programmed suicide” of infected mammal host cells. A recent reports documents that interferons molecule (type I,II and III) are little or not expressed in cells and tissues infected by SARS-CoV-2 whereas in contrast high level of chemokines and interleukin type 6, a pro inflammatory cytokine involved in innate immunity were observed [25]. The authors observed that the ongoing infection down regulated a large panel of genes among which are our genes making hybrid dsRNA. This puzzling transcriptomic picture regarding the classic immune responses to viral infection suggests a scenario for which SARS-CoV-2 induced molecular pathways in infected cells derailed from expected patterns. Recently, authors reports that they have retrieved the full catalogue of human proteins that bind with strong affinity to the 26 tagged SARS-CoV-2 proteins by double hybrid technology [26]. The datasets outlines that 42 human proteins are strong candidates for high affinity with viral partners (corrected p value from 1.41E-29 to 0.00578) and other 280 human proteins show weak statistical evidence for interaction [26]. Among this first group (n=42), 13 are mitochondrial proteins, each one interacting with different viral proteins and 6 are controsomal proteins interacting with a unique viral partner. Mitochondrial proteins are also heavily represented in the second group (n=280). In the full dataset of interacting human proteins, no protein linked to Dicer/ago pathways were found, which suggests that the RNAi process is not altered by the SARS-CoV-2 viral infection [26]. All together these information hint that the physiopathology landscape provoked by SARS-CoV-2 infection might be multifactorial including inactivation of major host proteins by high affinity binding with a viral partner along with drastic degradation of host messenger by RNAi machinerie.

## Method

### Viral sequences

We obtained the sequences of Severe acute respiratory syndrome coronavirus 2 (SARS-CoV-2); previously provisionally named 2019-nCoV, Middle East respiratory syndrome-related coronavirus (MERS-CoV), SARS coronavirus (SARS-CoV), Human coronavirus 229E (HCoV-229E) and Human coronavirus OC43 (HCoV-OC43) from NCBI GeneBank (RefSeq accession: NC_045512.2, NC_019843, NC_004718.3, NC_002645.1 and NC_006213.1) [1, 2, 27–30].

### Human transcriptome

RNA products annotated on the Genome Reference Consortium Human Build 38 patch release 13 (GRCh38.p13) were downloaded from NCBI FTP site (RefSeq accession: GCF_000001405.39).

### Identification of potential double-stranded RNA fragments

Our previous published analysis, along with other published works, confirmed that the catalogue of siRNAs is far more complex and extensive than previously thought and that it encompasses larger sets of the transcriptome [31–34]. A study has shown that an extensive presence of dsRNAs in the *Drosophila* and *C. elegans* obtained by high-throughput sequencing involves many categories of RNA including mRNAs in which miRNA and lncRNA populations appear a minority component [34]. Following the same approach as the one used by Pasquier et al. [31], each virus sequence have been cut up into short overlapping sequences of 15 bases. These 15-bases sequences were extracted at intervals of 6 bases to ensure that any sequence of at least 20 bases belonging to the complete virus sequence includes at least one of the 15-base sequences. The potential pairing were computed by aligning short 15-base sequences on RNA sequences with the STAR RNA-seq aligner [35] and by retaining only perfect matches for which the short sequence was reverse complemented. We then extended the matches to obtain the maximum alignment length. We subsequently applied a second post-processing step to eliminate duplicate alignments and to remove sequences aligned over less than 20 bases. This procedure has been applied to compute the potential pairing between RNA viruses and human transcripts and also to identify the possible internal pairing within RNA virus sequences. Briefly regarding SARS-CoV-2, the pairing segment was observed in all the isoforms (2 known and 13 predicted) of the human transcripts of USP30. The pairings were identified at the junction between exons 17 and 18 of DNAJC13, into exon 10 of ELP4, exon 12 of FBXO21, exon 2 of FLRT2, exon 13 of USP30, exon 16 of USP31 and exon 4 of ZNF443. The sequence of 20 bases, originating from orf3a of SARS-CoV-2 has the potential to target the sequence AAAGATAGAGAAAAGGGGCT which is shared by 15 predicted and/or found transcripts of USP30. On the other side, regarding USP30 gene, the data gathered by the “1000 genomes project” shows that this 20 bases sequence is present in all Homo Sapiens with the exception of 0,49 % of Han Chinese in Bejing that have a mutation A/T in 2nd position within this matching sequence [36]. Finally DNAJC13 is an endosome-related protein and believed to regulate endosomal membrane trafficking. Mutations provoke Parkinson disease and neurodegeneration [25]. In parallel the SARS-CoV-2 fragments that pair with USP31, FLRT2, FBXO21 were found in only 5, 4 and 5 isoforms (predicted or experimentally found) respectively. Regarding the MERS virus, the collection of hybrid dsRNA seems more disparate with the targeting of 2 ionic channels and a GTPase of Rho subfamily. The analysis of no virulent viruses (HCoV-229E and HCoV-OC43) did not reveal a targeting of essential genes except the coding genes: SSH1 (Protein tyrosine phosphatase involved in actin filament dynamics in cellular lamellipodia formation), PTP4A1 (Protein tyrosine phosphatase involved in progression G1-S mitosis), GCNA (protein with acidic domain involved in genome stability for which mutants accumulate DNA-protein crosslinks resulting in chromosome instability), COX11 (a component of the mitochondrial respiratory chain, that catalyzes the electron transfer from reduced cytochrome c to oxygen) and finally SRGAP1 (GTPase activator involved in neuronal migration) (see NCBI database for information).

### Identification of intra double-stranded RNA sequences in Coronaviridae

In parallel, we also investigate whether the structure of the 5 tested coronaviruses RNAs permits the formation of intermolecular double-stranded structures. We computed the potential internal pairing minimally over 20 bases length and identified only one candidate in SARS-CoV, involving a sequence of 22 bases starting at position 25961 and located in orf3a. This highlights that coronaviruses, under pressure of selection, have evolutionary succeeded in escaping the vigilance of RNAi host defense strategy by minimizing internal dsRNA structure.

### Assesment of the scope of hybrid duplexing against other species or viruses

To assess whether the sequences identified in SARS-CoV-2 could impact other species, we used BLAST to calculate the similarities between each human sequence and the following 12 other species: Gorilla gorilla, Pan troglodytes, Pongo abelii, Papio anubis, Macaca mulatta, Sapajus apella, Mus musculus, Rattus norvegicus, Pteropus vampyrus, Sus scrofa, Bos taurus and Felis catus. The potentiality of hybrid pairing depends on the species and on candidate mRNAs in the host (figure 1 and supplementary tables S2 to S8). USP30 features the only human-specific hybrid pairing. Conversely, using BLAST, we also calculated similarities of sequences originating from SARS-CoV-2 with the following viruses: Bat coronavirus RaTG13, SARS-CoV, MERS-CoV, HcoV-229E and HcoV-OC43. The results are presented in figure 2 and supplementary tables S9 to S15).

## Acknowledgments

This work was supported by the ANR grant “Methylclonome” ANR-12-BSV6-006-01 to Alain Robichon and Claude Pasquier. This work was also supported by the French National Research Agency (ANR) through the LABEX SIGNALIFE program (reference # ANR-11-LABX-0028-01).

## Author contributions

C.P. did the bio-informatic analysis. C.P. and A.R. conceive the conceptual framework of this study and wrote the paper.

---

**Supplemental Table S1** Hybrid dsRNAs between human transcriptome and SARS-CoV-2 RNA. The table details hybrid duplexing RNA between SARS-CoV-2 and humans. The analysis detailed all human transcript isoforms for each targeted gene by the viral RNA. Four other coronaviruses were analyzed in comparison with SARS-CoV-2.

**Supplemental Tables S2 to S15** Alignments of human and SARS-CoV-2 strands within the hybrid dsRNAs collection with respectively other mammal and virus species. The tables represent the alignments of the identified 20 bases length sequences between different mammal species and between different coronaviruses. These sequences correspond to the 7 genes identified by computational search as potentially generating hybrid dsRNA between SARS-CoV-2 and humans. The alignment analysis was conducted with the human strand of hybrid dsRNA regarding the similarities with other mammal species. The viral strand regarding similarities with other coronaviruses was compared in parallel.

