## Supplemental data-Tables S2-S15 for "Computational search of hybrid human/ SARS-CoV-2 dsRNA reveals unique viral sequences that diverge from those of other coronavirus strains"

| Specie | Common name | Sequence in host | Gene in host | % sim. |
| --- | --- | --- | --- | --- |
| Homo sapiens | human | AGACAGCTAAGTAGACATTT | DNAJC13 | 100 |
| Gorilla gorilla | western gorilla | AGACAGCTAAGTAGACATTT | DNAJC13 | 100 |
| Pan troglodytes | chimpanzee | AGACAGCTAAGTAGACATTT | DNAJC13 | 100 |
| Pongo abelii | sumatran orangutan | AGACAGCTAAGTAGACATTT | DNAJC13 | 100 |
| Papio anubis | olive baboon | AGACAGCTAAGTAGACATTT | DNAJC13 | 100 |
| Macaca mulatta | rhesus monkey | AGACAGCTAAGTAGACATTT | DNAJC13 | 100 |
| Sapajus apella | tufted capuchin | AGACAGCTAAGTAGACATTT | DNAJC13 | 100 |
| Mus musculus | house mouse | AGACAGCTAAGTAGACATTT | DNAJC13 | 90 |
| Rattus norvegicus | norway rat | AGACAGCTAAGTAGACATTT | DNAJC13 | 90 |
| Pteropus vampyrus | large flying fox | AGACAGCTAAGTAGACATTT | DNAJC13 | 100 |
| Sus scrofa | pig | AGACAGCTAAGTAGACATTT | DNAJC13 | 100 |
| Bos taurus | cattle | AGACAGCTAAGTAGACATTT | DNAJC13 | 100 |
| Felis catus | domestic cat | AGACAGCTAAGTAGACATTT | DNAJC13 | 100 |

**S2 Presence in other species of the 20-bases sequence of DNAJC13 theoretically capable of forming dsRNA with SARS-CoV-2**

| Specie | Common name | Sequence in host | Gene in host | % sim. |
| --- | --- | --- | --- | --- |
| Homo sapiens | human | TTCTGCATGTGCAAGCATTT | ELP4 | 100 |
| Gorilla gorilla | western gorilla | TTCTGCATGTGCAAGCATTT | ELP4 | 45 |
| Pan troglodytes | chimpanzee | TTCTGCATGTGCAAGCATTT | ELP4 | 45 |
| Pongo abelii | sumatran orangutan | TTCTGCATGTGCAAGCATTT | ELP4 | 45 |
| Papio anubis | olive baboon | TTCTGCATGTGCAAGCATTT | ELP4 | 45 |
| Macaca mulatta | rhesus monkey | TTCTGCATGTGCAAGCATTT | ELP4 | 45 |
| Sapajus apella | tufted capuchin | TTCTGCATGTGCAAGCATTT | ELP4 | 100 |
| Mus musculus | house mouse | TTCTGCATGTGCAAGCATTT | ELP4 | 40 |
| Rattus norvegicus | norway rat | TTCTGCATGTGCAAGCATTT | ELP4 | 35 |
| Pteropus vampyrus | large flying fox | TTCTGCATGTGCAAGCATTT | ELP4 | 45 |
| Sus scrofa | pig | TTCTGCATGTGCAAGCATTT | ELP4 | 45 |
| Bos taurus | cattle | TTCTGCATGTGCAAGCATTT | ELP4 | 35 |
| Felis catus | domestic cat | TTCTGCATGTGCAAGCATTT | ELP4 | 40 |

**S3 Presence in other species of the 20-bases sequence of ELP4 theoretically capable of forming dsRNA with SARS-CoV-2**

| Specie | Common name | Sequence in host | Gene in host | % sim. |
| --- | --- | --- | --- | --- |
| Homo sapiens | human | AGAGAGAGAATGTCTTTCAT | FBXO21 | 100 |
| Gorilla gorilla | western gorilla | AGAGAGAGAATGTCTTTCAT | FBXO21 | 100 |
| Pan troglodytes | chimpanzee | AGAGAGAGAATGTCTTTCAT | FBXO21 | 100 |
| Pongo abelii | sumatran orangutan | AGAGAGAGAATGTCTTTCAT | FBXO21 | 90 |
| Papio anubis | olive baboon | AGAGAGAGAATGTCTTTCAT | FBXO21 | 80 |
| Macaca mulatta | rhesus monkey | AGAGAGAGAATGTCTTTCAT | FBXO21 | 80 |
| Sapajus apella | tufted capuchin | AGAGAGAGAATGCCTTTCAT | FBXO21 | 75 |
| Mus musculus | house mouse | AGAGAGAGAATGTCTTTCAT | FBXO21 | 40 |
| Rattus norvegicus | norway rat | AGAGAGAGAATGTCTTTCAT | FBXO21 | 40 |
| Pteropus vampyrus | large flying fox | AGAGAGAGAATGTCTTTCAT | FBXO21 | 35 |
| Sus scrofa | pig | AGAGAGAGAATGTCTTTCAT | FBXO21 | 35 |
| Bos taurus | cattle | AGAGAGAGAATGTCTTTCAT | FBXO21 | 45 |
| Felis catus | domestic cat | AGAGAGAGAATGTCTTTCAT | FBXO21 | 45 |

**S4 Presence in other species of the 20-bases sequence of FBXO21 theoretically capable of forming dsRNA with SARS-CoV-2**

| Specie | Common name | Sequence in host | Gene in host | % sim. |
| --- | --- | --- | --- | --- |
| Homo sapiens | human | CATCTATGCTATTCTTGGGT | FLRT2 | 100 |
| Gorilla gorilla | western gorilla | CATCTATGCTATTCTTGGGT | FLRT2 | 45 |
| Pan troglodytes | chimpanzee | CATCTATGCTATTCTTGGGT | FLRT2 | 100 |
| Pongo abelii | sumatran orangutan | CATCTATGCTATTCTTGGGT | FLRT2 | 45 |
| Papio anubis | olive baboon | CATCCATGCTATTCTTGGGT | FLRT2 | 85 |
| Macaca mulatta | rhesus monkey | CATCTATGCTATTCTTGGGT | FLRT2 | 45 |
| Sapajus apella | tufted capuchin | CATCTATGCTATTCTTGGGT | FLRT2 | 75 |
| Mus musculus | house mouse | CATCTATGCTATTCTTGGGT | FLRT2 | 45 |
| Rattus norvegicus | norway rat | CATCTATGCTATTCTTGGGT | FLRT2 | 45 |
| Pteropus vampyrus | large flying fox | CATCTATGCCATTCTTGGGT | FLRT2 | 60 |
| Sus scrofa | pig | CATCTATGCTATTCTTGGGT | FLRT2 | 45 |
| Bos taurus | cattle | CATCTATGCTATTCTTGGGT | FLRT2 | 40 |
| Felis catus | domestic cat | CATCTATGCTATTCTTGGGT | FLRT2 | 45 |

**S5 Presence in other species of the 20-bases sequence of FLRT2 theoretically capable of forming dsRNA with SARS-CoV-2**

| Specie | Common name | Sequence in host | Gene in host | % sim. |
| --- | --- | --- | --- | --- |
| Homo sapiens | human | GCACAAAATAATCACCAACA | USP31 | 100 |
| Gorilla gorilla | western gorilla | GCACAAAATAATCACCAACA | USP31 | 100 |
| Pan troglodytes | chimpanzee | GCACAAAATAATCACCAACA | USP31 | 100 |
| Pongo abelii | sumatran orangutan | GCACAAAATAATCACCAACA | USP31 | 35 |
| Papio anubis | olive baboon | GCACAAAATAATCACCAACA | USP31 | 85 |
| Macaca mulatta | rhesus monkey | GCACAAAATAATCACCAACA | USP31 | 85 |
| Sapajus apella | tufted capuchin | GCACAAAATAATCACCAACA | USP31 | 100 |
| Mus musculus | house mouse | GCACAAAATAATCACCAACA | USP31 | 45 |
| Rattus norvegicus | norway rat | GCACAAAATAATCACCAACA | USP31 | 40 |
| Pteropus vampyrus | large flying fox | GCACAAAATAATCACCAACA | USP31 | 35 |
| Sus scrofa | pig | GCACAAAATAATCACCAACA | USP31 | 40 |
| Bos taurus | cattle | GCACAAAATAATCACCAACA | USP31 | 55 |
| Felis catus | domestic cat | GCACAAAATAATCACCAACA | USP31 | 40 |

**S6 Presence in other species of the 20-bases sequence of USP31 theoretically capable of forming dsRNA with SARS-CoV-2**

| Specie | Common name | Sequence in host | Gene in host | % sim. |
| --- | --- | --- | --- | --- |
| Homo sapiens | human | GAAACCATATGATTGTAAAG | ZNF443 | 100 |
| Gorilla gorilla | western gorilla | GAAACCATATGATTGTAAAG | ZNF443 | 100 |
| Pan troglodytes | chimpanzee | GAAACCATATGATTGTAAAG | ZNF443 | 100 |
| Pongo abelii | sumatran orangutan | GAAACCATATGATTGTAAAG | ZNF443 | 100 |
| Papio anubis | olive baboon | GAAACCATATGATTGTAAAG |  |  |
| Macaca mulatta | rhesus monkey | GAAACCATATGATTGTAAAG |  |  |
| Sapajus apella | tufted capuchin | GAAACCATATGATTGTAAAG |  |  |
| Mus musculus | house mouse | GAAACCATATGATTGTAAAG | Zfp709 | 45 |
| Rattus norvegicus | norway rat | GAAACCATATGATTGTAAAG | Zfp709 | 40 |
| Pteropus vampyrus | large flying fox | GAAACCATATGATTGTAAAG |  |  |
| Sus scrofa | pig | GAAACCATATGATTGTAAAG |  |  |
| Bos taurus | cattle | GAAACCATATGATTGTAAAG |  |  |
| Felis catus | domestic cat | GAAACCATATGATTGTAAAG |  |  |

**S7 Presence in other species of the 20-bases sequence of ZNF443 theoretically capable of forming dsRNA with SARS-CoV-2. The table lists only the orthologs of ZNF443 described in the database Homologene at NCBI**

| Specie | Common name | Sequence in host | Gene in host | % sim. |
| --- | --- | --- | --- | --- |
| Homo sapiens | human | AAAGATAGAGAAAAGGGGCT | USP30 | 100 |
| Gorilla gorilla | western gorilla | AAAGATGGAGAAAAGGGGCT | USP30 | 95 |
| Pan troglodytes | chimpanzee | AAAGATGGAGAAAAGGGGCT | USP30 | 95 |
| Pongo abelii | sumatran orangutan | AAAGATGGAGAAAAGGGGCT | USP30 | 95 |
| Papio anubis | olive baboon | AAAGATGGAGAAAAGCGGCT | USP30 | 90 |
| Macaca mulatta | rhesus monkey | AAAGATGGAGAAAAGCGGCT | USP30 | 90 |
| Sapajus apella | tufted capuchin | AAAGATGGAGAAAAGGGGCT | USP30 | 75 |
| Mus musculus | house mouse | AAAGATAGAGAAAAGGGGCT | USP30 | 50 |
| Rattus norvegicus | norway rat | AAAGATAGAGAAAAGGGGCT | USP30 | 50 |
| Pteropus vampyrus | large flying fox | AAAGATAGAGAAAAGGGGCT | USP30 | 50 |
| Sus scrofa | pig | AAAGATAGAGAAAAGGGGCT | USP30 | 50 |
| Bos taurus | cattle | AAAGATAGAGAAAAGGGGCT | USP30 | 45 |
| Felis catus | domestic cat | AAAGATAGAGAAAAGGGGCT | USP30 | 45 |

**S8 Presence in other species of the 20-bases sequence of USP30 theoretically capable of forming dsRNA with SARS-CoV-2**

| Virus name | Virus sequence | % sim. |
| --- | --- | --- |
| SARS-CoV-2 | AAATGTCTACTTAGCTGTCT | 100 |
| Bat coronavirus RaTG13 | AAATGTCTACCTAGCTGTCT | 95 |
| SARS-CoV | AAATGTCTACTTAGCTGTCT | 45 |
| MERS-CoV | AAATGTCTACTTAGCTGTCT | 45 |
| HCoV-229E | AAATGTCTACTTAGCTGTCT | 45 |
| HCoV-OC43 | AAATGTCTACTTAGCTGTCT | 45 |

**S9 Similarities in other viruses of the 20-bases sequence of SARS-CoV-2 theoretically capable of forming dsRNA with human gene DNAJC13**

| Virus name | Virus sequence | % sim. |
| --- | --- | --- |
| SARS-CoV-2 | AAATGCTTGCACATGCAGAA | 100 |
| Bat coronavirus RaTG13 | AAATGCTTGCACATGCAGAA | 95 |
| SARS-CoV | AAATGCTTGCTCATGCAGAA | 75 |
| MERS-CoV | AAATACTTGCACATGCAGAA | 65 |
| HCoV-229E | AAATGGTTGCACATGCAGAA | 65 |
| HCoV-OC43 | AAATGCTTGCACATGCAGAA | 45 |

**S10 Similarities in other viruses of the 20-bases sequence of SARS-CoV-2 theoretically capable of forming dsRNA with human gene ELP4**

| Virus name | Virus sequence | % sim. |
| --- | --- | --- |
| SARS-CoV-2 | ATGAAAGACATTCTCTCTCT | 100 |
| Bat coronavirus RaTG13 | ATGAAAGACATTCTCTCTCT | 100 |
| SARS-CoV | ATGAGAGACATCCTCTCTCT | 85 |
| MERS-CoV | ATGAAAGACATTCTCTCTCT | 45 |
| HCoV-229E | ATGAAAGACATTCTCTCTCT | 45 |
| HCoV-OC43 | ATGAAAGACATTCTCTCTCT | 60 |

**S11 Similarities in other viruses of the 20-bases sequence of SARS-CoV-2 theoretically capable of forming dsRNA with human gene FBXO21**

| Virus name | Virus sequence | % sim. |
| --- | --- | --- |
| SARS-CoV-2 | ACCCAAGAATAGCATAGATG | 100 |
| Bat coronavirus RaTG13 | ACCCAAGAATAGCATAGATG | 80 |
| SARS-CoV | ACCCAAGAATAGCATAGATG | 40 |
| MERS-CoV | ACCCAAGAATAGCATAGATG | 50 |
| HCoV-229E | ACCCAAGAATAGCATAGATG | 40 |
| HCoV-OC43 | ACCCAAGAATAGCATAGATG | 40 |

**S12 Similarities in other viruses of the 20-bases sequence of SARS-CoV-2 theoretically capable of forming dsRNA with human gene FLRT2**

| Virus name | Virus sequence | % sim. |
| --- | --- | --- |
| SARS-CoV-2 | TGTTGGTGATTATTTTGTGC | 100 |
| Bat coronavirus RaTG13 | TGTTGGTGACTATTTTGTGC | 95 |
| SARS-CoV | TGTTGGTGATTACTTTGTGC | 90 |
| MERS-CoV | TGTTGGTGATTATTTTGTGC | 55 |
| HCoV-229E | TGTTGGTGATTATTTTGTGC | 60 |
| HCoV-OC43 | TGTTGGTGATTATTTTGTGC | 65 |

**S13 Similarities in other viruses of the 20-bases sequence of SARS-CoV-2 theoretically capable of forming dsRNA with human gene USP31**

| Virus name | Virus sequence | % sim. |
| --- | --- | --- |
| SARS-CoV-2 | CTTTACAATCATATGGTTTC | 100 |
| Bat coronavirus RaTG13 | CTTTACAATCATATGGTTTC | 45 |
| SARS-CoV | CTTTACCATCATATGGTTTC | 70 |
| MERS-CoV | CTTTACAATCATATGGTTTC | 65 |
| HCoV-229E | CTTTACAATCATATGGTTTC | 45 |
| HCoV-OC43 | CTTTACAATCATATGGTTTC | 60 |

**S14 Similarities in other viruses of the 20-bases sequence of SARS-CoV-2 theoretically capable of forming dsRNA with human gene ZNF443**

| Virus name | Virus sequence | % sim. |
| --- | --- | --- |
| SARS-CoV-2 | AGCCCCTTTTCTCTATCTTT | 100 |
| Bat coronavirus RaTG13 | AGCCCCTTTTCTCTATCTTT | 45 |
| SARS-CoV | AGCCCCATTTCTTTATCTTT | 90 |
| MERS-CoV | AGCCCCTTTTCTCTATCTTT | 45 |
| HCoV-229E | AGCCCCTTTTCTCTATCTTT | 45 |
| HCoV-OC43 | AGCCCCTTTTCTCTATCTTT | 40 |

**S15 Similarities in other viruses of the 20-bases sequence of SARS-CoV-2 theoretically capable of forming dsRNA with human gene USP30**
